# Accurate DNA Methylation Predictor for *C9orf72* Repeat Expansion Alleles in the Pathogenic Range

**DOI:** 10.1101/2025.03.20.643775

**Authors:** Naren Ramesh, Alexandria Evans, Kevin Wojta, Zhongan Yang, Marco MP Boks, René S. Kahn, Sterre C.M. de Boer, Sven J. van der Lee, Yolande A.L. Pijnenburg, Lianne M. Reus, Roel A. Ophoff

## Abstract

The hexanucleotide (G_4_C_2_) repeat expansion in the promoter region of *C9orf72* is the most frequent genetic cause of frontotemporal dementia (FTD) and amyotrophic lateral sclerosis (ALS). In this study, we conducted a genome-wide DNA methylation (DNAm) analysis using EPIC version 2 (EPICv2) arrays on an FTD cohort comprising 27 carriers and 250 non-carriers of the pathogenic *C9orf72* repeat expansion from the Amsterdam Dementia Cohort.

We identified differentially methylated CpGs probes associated with the pathogenic *C9orf72* expansion and used these findings to create a DNAm Least Absolute Shrinkage and Selection Operator (LASSO) predictor to identify repeat expansion carriers. Eight CpG sites at the *C9orf72* locus were significantly differentially hypermethylated in repeat expansion carriers compared to non-carriers. The LASSO model predicted repeat expansion status with an average accuracy of 98.6%. The LASSO predictor was further validated in an independent cohort of 2,548 subjects with available EPICv2 data, identifying four *C9orf72* repeat expansion carriers, subsequently confirmed by repeat-primed PCR. This result not only illustrates the accuracy of the DNAm predictor of *C9orf72* repeat expansion carriers but also suggests that repeat expansion carriers may be more prevalent than expected. The identification of a highly accurate DNAm biomarker for a repeat expansion locus associated with neurodegenerative disorders may provide great value for studying this locus. The approach holds significant promise for investigating this and other repeat expansion loci, particularly given the growing interest in epigenetic epidemiological studies involving large cohorts with available DNAm data.

**Graphical abstract (*optional*):** 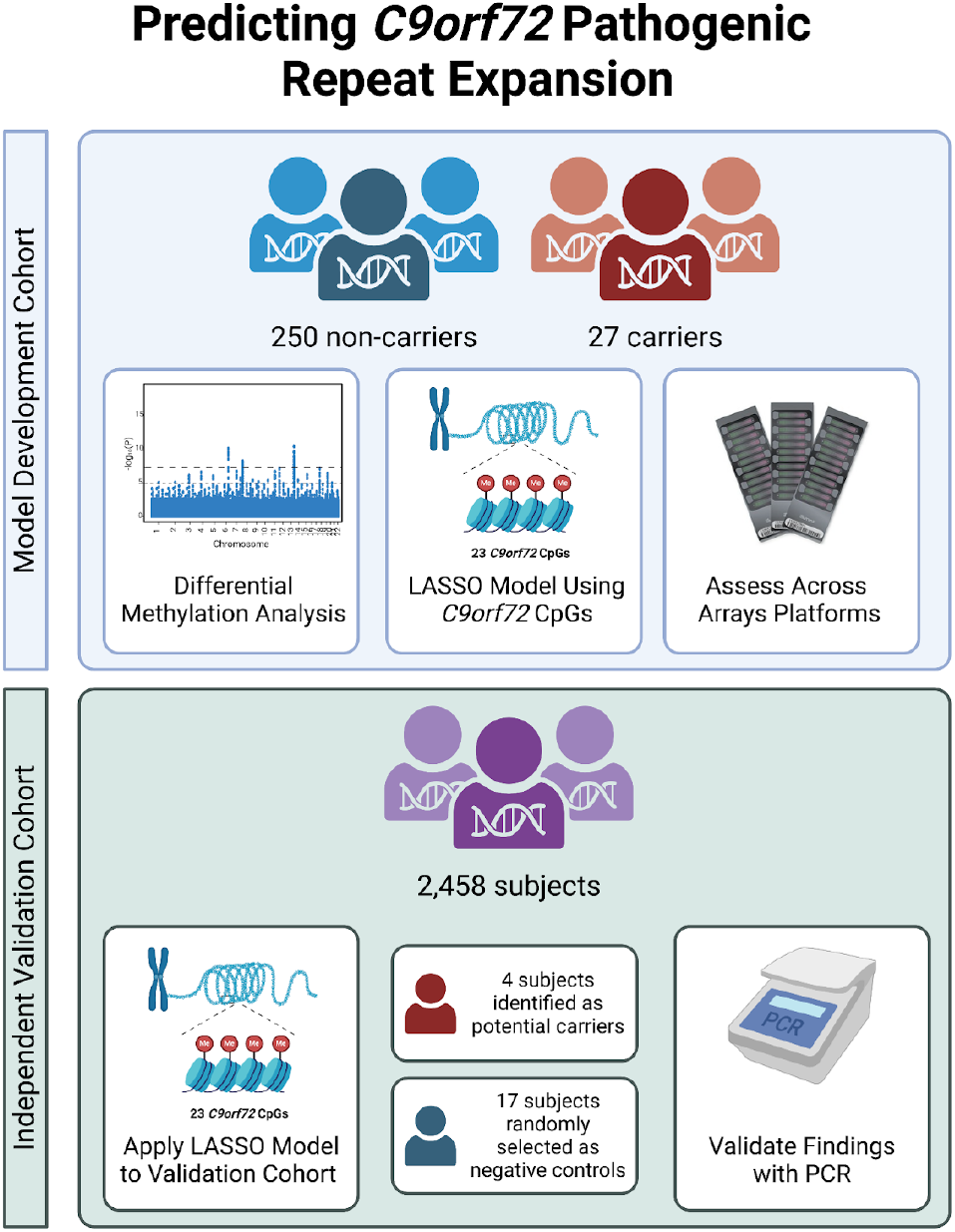

## Introduction

An increased hexanucleotide repeat expansion of GGGGCC (G_4_C_2_) in the noncoding region of the *C9orf72* gene is one of the most common genetic causes for amyotrophic lateral sclerosis (ALS) and frontotemporal dementia (FTD)^1–3^. This repeat expansion is found in 6-7% of ALS/FTD without a family history for the disease, 39.3% of familial ALS cases, and 24.8% of familial FTD cases among European ancestry patients^4^. While the precise cut-off for non-pathogenic and pathogenic repeat lengths is debated, expansions greater than 30 repeats are generally considered to be pathogenic for ALS and FTD, with some affected patients having expansions into the thousands of repeat units^5–7^.

Identifying pathogenic *C9orf72* repeat expansions at both the individual and population levels remains crucial. Clinical symptoms associated with *C9orf72* repeat expansion are serious and diverse, with neurodegenerative symptoms manifesting in ALS, FTD, and movement disorders such as Parkinsonism but also resulting in behavioral phenotypes observed in FTD and psychiatric-related phenotypes such as schizophrenia and bipolar disorder^8,9^. *C9orf72* expansion is usually tested after suspicion for FTD or ALS or a positive family history for the mutation, but patients with other disorders and symptoms are not examined for this mutation as part of a standard diagnostic routine^10^. FTD, especially behavioral variant (bvFTD), is often difficult to diagnose or identify due to the heterogeneity of clinical symptoms of the disease, diagnostic overlap with other neurodegenerative disorders, similarity with psychiatric phenotypes, and the lack of biomarkers^11,12^. High percentages of patients with bvFTD, ranging from 50-70%, are misdiagnosed as having other neuropsychiatric conditions or dementia^13–15^ causing an average six years delay between symptom onset and the diagnosis of bvFTD^16^.

Standard screening for the *C9orf72* repeat expansion is performed using repeat-primed PCR^2^ or testing with Southern blotting^17^. Southern blotting allows for more precise assessment of the length of the repeat expansion, but the approach is very labor intensive, hard to scale, and requires large amounts of genomic DNA. PCR-based methods, on the other hand, are well-established for the detection of expanded alleles, can be used in higher-throughput settings, but are not suitable for estimating the actual repeat length itself. More recently, computational methods for analysis of whole-genome sequence data, such as ExpansionHunter^18^, have become available for the detection of repeat expansions but the availability of sequence data remains a rate-limiting step. In the near future, we expect that technological advances with long read sequencing will further advance molecular assessment of *C9orf72* repeat expansion alleles^19^.

Epigenetic features provide valuable insights into the mechanisms underlying *C9orf72* pathogenesis and may help address challenges related to identifying *C9orf72* expansions. Notably, studies have shown that patients with FTD and ALS often exhibit hypermethylation in the *C9orf72* gene promoter and within the repeat expansion itself ^20,21^. Building on these findings, we conducted a genome-wide analysis of differential DNA methylation profiles in FTD patients from the Amsterdam Dementia Cohort (ADC) with both pathogenic and non-pathogenic *C9orf72* repeat alleles to assess the extent of DNA methylation changes associated with these expansions. We identified significant evidence for epigenetic *cis* effects of the repeat expansion and used this insight to develop an accurate DNAm predictor for the detection of pathogenic repeat expansion alleles at this locus.

## Results

### Sample characteristics

Demographic and clinical characteristics of the ADC cohort are presented in **Table S1**. For the remainder of this study, we will be referencing expansions that are larger than 45 repeat as pathogenic repeat expansions. This cohort included n = 27 FTD patients with a known pathogenic *C9orf72* repeat expansion (mean age of 62.2 [6.7], 37.0% female) and n = 250 FTD patients without a pathogenic *C9orf72* repeat expansion (mean age of 63.3 [8.4], 37.6% female). Carrier status of the pathogenic *C9orf72* hexanucleotide repeat expansion was established as part of the diagnostic assessment performed at the Alzheimer Center Amsterdam. Pathological *C9orf72* repeat length carriers did not differ from the non-carriers regardless of diagnosis status in terms of age (*p* = 1.00), sex (*p* = 1.00), and epigenetic smoking score (*p* = 1.00)^22^. Based on imputed cell composition^23^, pathological *C9orf72* repeat expansion carriers did not differ from the non-carriers in terms of CD8+ T cells (*p* = 0.51), CD4+ T cells (*p* = 0.52), natural killer cells (*p* = 0.45), B cells (*p* = 0.60), monocytes (*p* = 1.00), and neutrophils (p= 0.14).

### Genome-Wide Differential Methylation Analysis of *C9orf72* Expansion Status

To identify the epigenetic signature of pathological *C9orf72* repeat lengths, we conducted a genome-wide differential methylation analysis on individuals with and without a pathological *C9orf72* repeat expansion, correcting for age, sex, cellular composition, epigenetic smoking score, and experimental batch (n=277 subjects, 837,312 CpG probes each). We identified eight CpG probes that exhibit differential methylation patterns between individuals with and without pathological *C9orf72* repeat lengths that exceed the genome-wide significance threshold (*p* = 4.4e-10) (**Table 1**; **Figure 1a**). This significance threshold was determined following a permutation analysis randomly shuffling expansion carrier status (of 27 subjects) 100 times. All eight CpGs were hypermethylated in the pathological repeat expansion carriers (**Figure 1d; Figure S2**). We did not find any evidence for genomic inflation of the genome-wide signal (**λ**_gc_= 1.006) suggesting that the observed associations are not due to systematic biases or confounding factors (**Figure 1b**). Notably, these eight most significantly differentially methylated probes were all located within the *C9orf72* gene region (**Figure 1e**). These associations highlight the potential biological (and/or clinical) relevance of methylation changes at this locus and may offer utility in identifying pathological *C9orf72* repeat expansion carriers.

**Figure 1.**
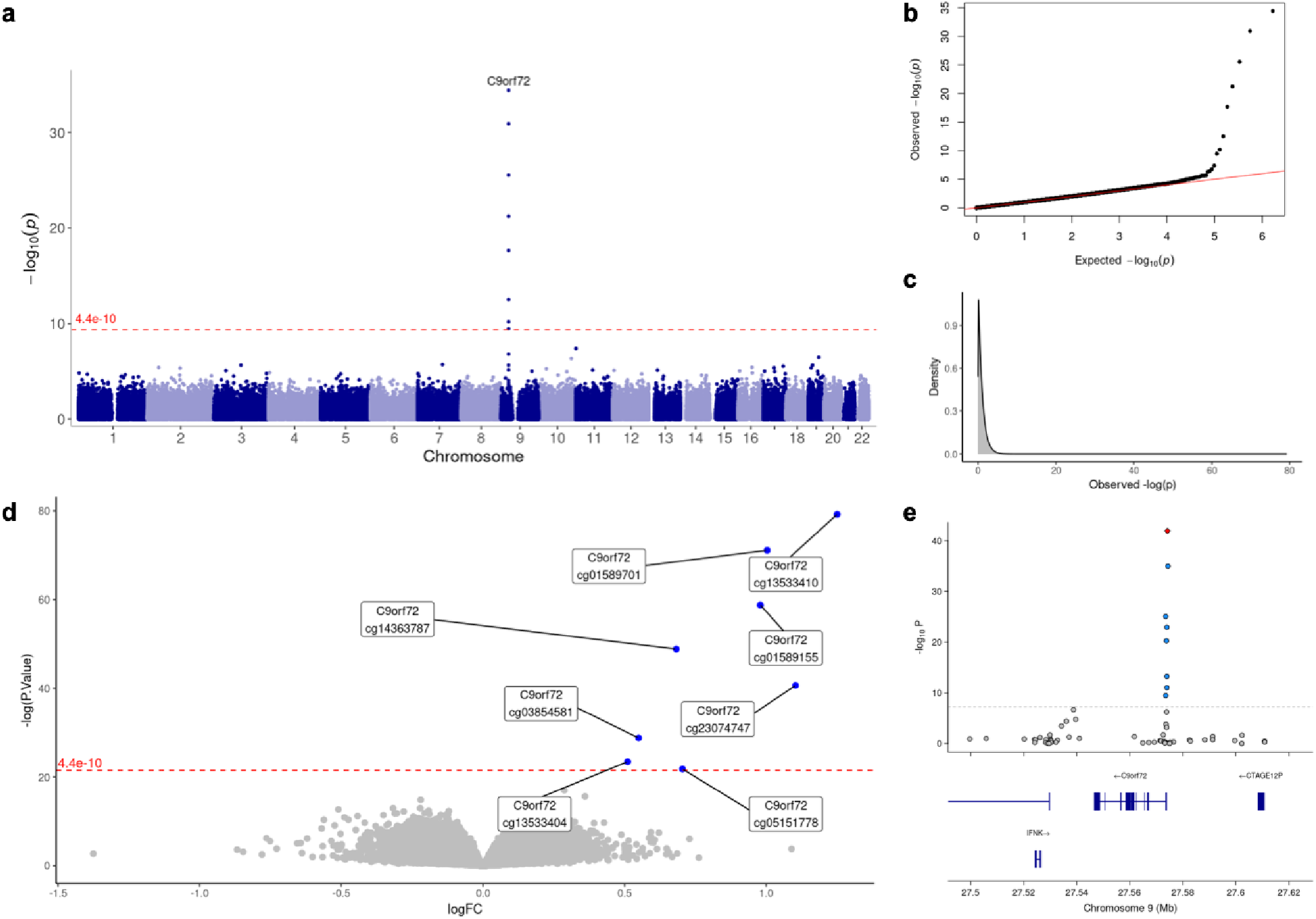
Hypermethylation in the *C9orf72* Gene Associated with Pathological *C9orf72* Repeat Expansion Carriers. **(a)** Manhattan plot showing *p*-values for CpGs tested for differential methylation patterns between carriers and non-carriers of *C9orf72* repeat expansions in the pathogenic range. The genome-wide significance threshold derived from a permutation analysis (*p =* 4.4e-10) is marked by the red dashed line. **(b)** QQ-plot showing the p value distribution and inflation (λ_gc_ value). **(c)** Density plot showing the observed p-value distribution. **(d)** Volcano plot showing statistical significance of DMA results against logFC illustrating direction of significance. Blue dots represent significant CpGs that are hypermethylated in carriers of *C9orf72* pathogenic repeat expansions. The genome-wide significance threshold derived from a permutation analysis (*p =* 4.4e-10) is marked by the red dashed line. **(e)** Locus Zoom Plot of Chromosome 9 between 27.5 Mb to 27.62 Mb including genes in the area. Blue dots indicate significantly differentially methylated CpG probes and the red diamond indicates the most statistically significant CpG.

**Table 1.**
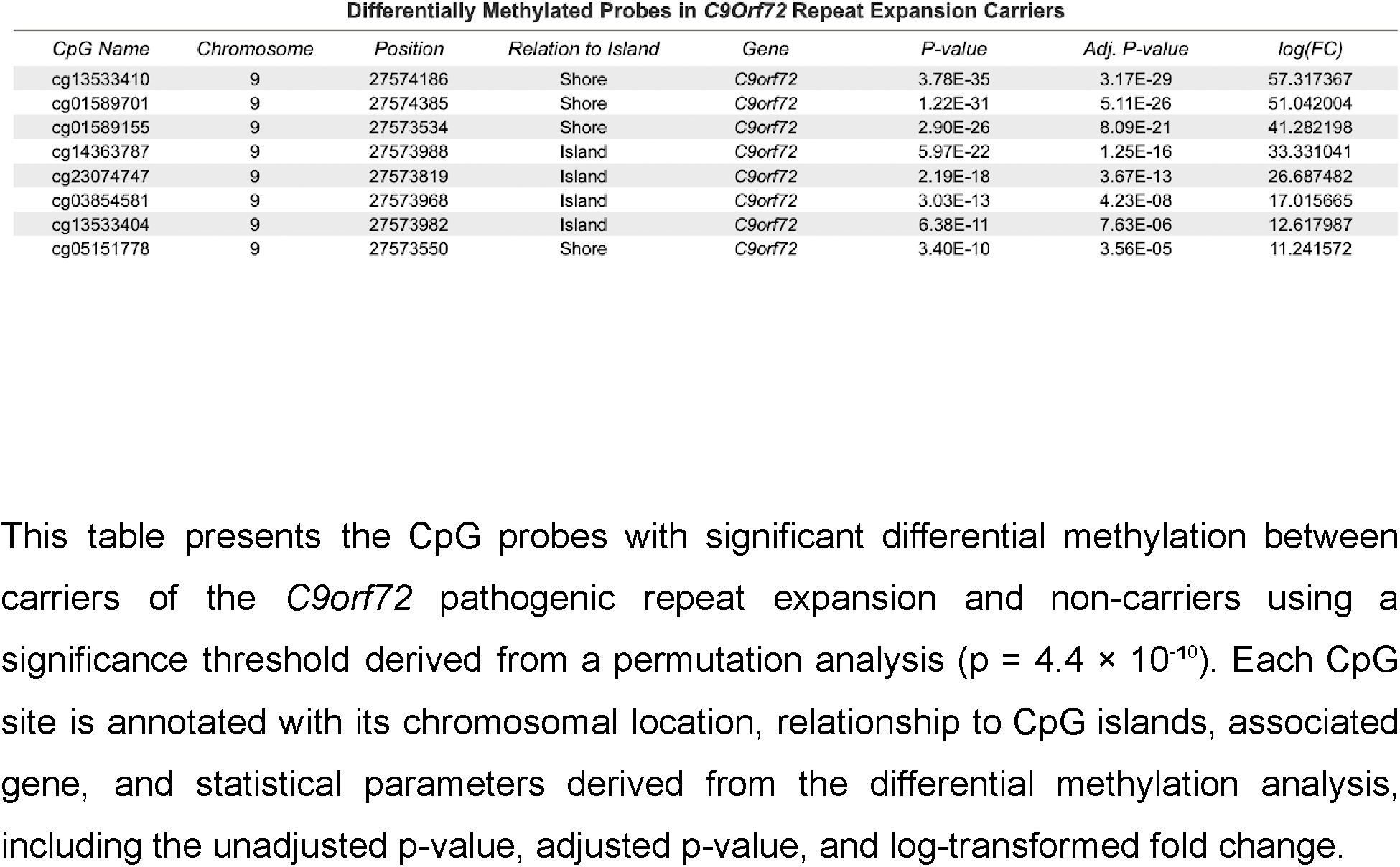
Differentially Methylated CpG Probes in *C9orf72* Pathogenic Repeat Expansion Carriers.

### Prediction of Expansion using CpGs in the *C9orf72* Region

We next examined whether we could predict pathological *C9of72* repeat lengths using methylation patterns of the CpG probes within the *C9orf72* gene. Given differential methylation patterns in probes within the *C9orf72* gene, we specified our analysis to 23 CpG probes that were found 1 kilobase upstream, 1 kilobase downstream, and within *C9orf72* itself. Principal components (PC) 1 and 2 on these *C9orf72* probes showed separation between individuals with and without pathological *C9orf72* repeat lengths (**Figure S3**). We further applied Least Absolute Shrinkage and Selection Operator (LASSO) regression to the 23 CpG probes to predict pathological *C9orf72* repeat lengths. The data was divided into a training set (70%) and a test set (30%), and we employed 10-fold cross-validation to optimize the model parameters and prevent model overfitting (**Figure S4b and S4c**). In the given test and train split, the LASSO regression model achieved 100% accuracy in predicting pathological *C9orf72* repeat expansion status on the test set (**Figure S4a**). The model selected 9 CpG probes as important predictors (**Table S2**). Five of these probes were found to be significantly differentially methylated between repeat carriers and non-carriers in our initial differential methylation analysis.

### Prediction Accuracy is Dependent on Array Technology

Next, we aimed to assess if predicting pathological *C9orf72* repeat expansion could be successful in previous generation Illumina DNA methylation array types limiting the LASSO model to CpGs present in EPICv1, Methyl450K, and Methyl 27K. Of the initial 23 CpG probes from EPICv2 within the *C9orf72* gene and 1 kb flanking it, 12 remained in EPICv1, 5 in Methyl450K, and 2 in Methyl27K (**Table 2**). Using the probes from each platform to predict pathogenic *C9orf72* repeat expansion, we found that the average accuracies were similar between EPICv2 (98.6%) and EPICv1 (99.0%) (*p* = 0.059). However, Methyl450K (94.1%) and Methyl27K (91.7%) were significantly less accurate compared to EPICv2 and EPICv1 (all *p* < 1e-15). (**Figure 2a**) Average Type I error rates were statistically similar EPICv2 (0.1%), EPICv1 (0.09%), and Methyl450K (0.2%) (all *p* > 0.1). However, Type I error rates were significantly higher for Methyl27K (0.7%) compared to Methyl450K (*p* = 2.5e-07), EPICv1 (*p* = 2.1e-10), and EPICv2 (*p* = 1.9e-09). (**Figure 2b**) Average Type II error rates were only nominally different between EPICv2 (11.4%) and EPICv1 (*p* = 0.028). Methyl450K (66.6%) and Methyl27K (75%) had significantly higher average Type II. (**Figure 2c)** Predicting pathological expansion status using CpG probes from EPICv2 and EPICv1 yielded high accuracy with low false positive and false negative rates. However, using probes from Methyl450K resulted in markedly lower accuracy with a significantly elevated false negative rate. The Methyl27K array had the lowest accuracy, with significantly elevated false positive and false negative rates.

**Figure 2.**
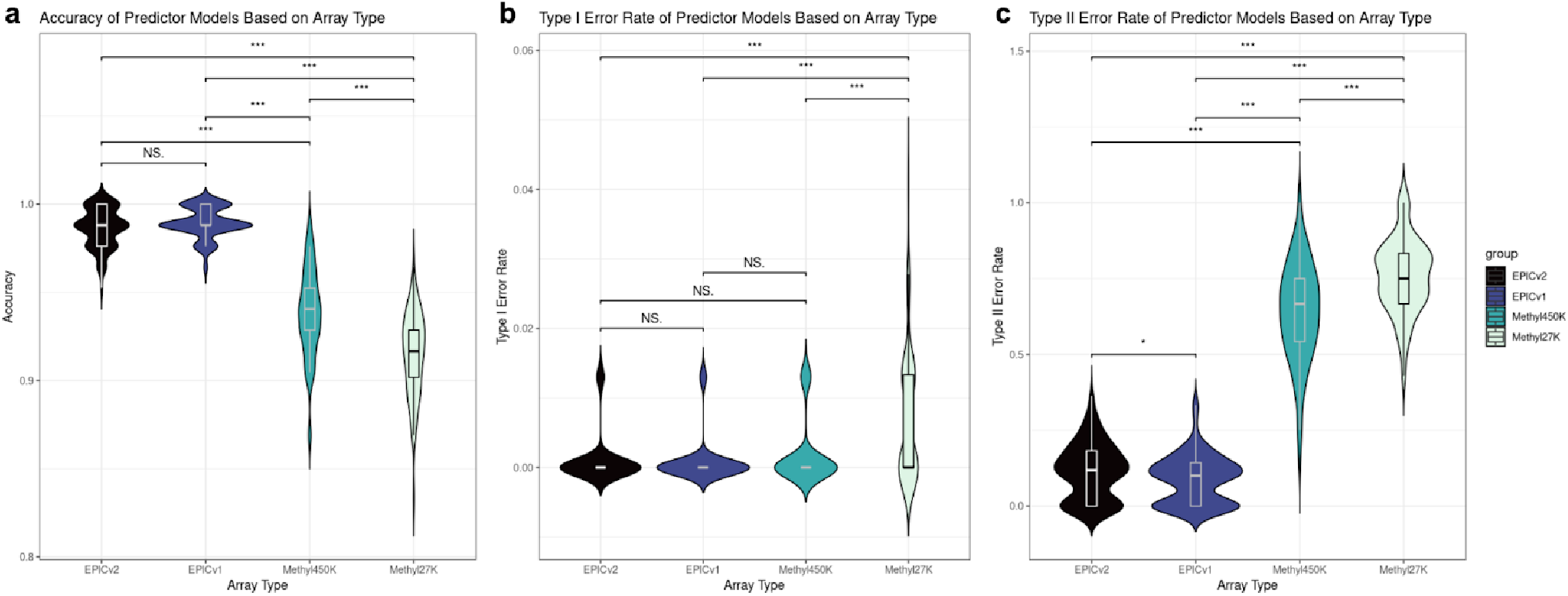
Accuracy, Type I Error, and Type II Error Across Array Type. Violin plots showing the distribution of **(a)** accuracy, **(b)** Type I error (false positive) rates, and **(c)** Type II error (false negative) rates from 100 iterations of training and testing a LASSO model using EPICv2 CpGs present in EPICv2 (23), EPICv1 (12), Methyl450K (5), and Methyl27K (2).

**Table 2.** CpG Probes in *C9orf72* Gene Region According to Platform.

| *Chr9 Position* | *EPICv2* | *EPICv1* | *Methyl450K* | *Methyl27K* |
| --- | --- | --- | --- | --- |
| 27561767 | cg13533245 | - | - | - |
| 27564906 | cg01126010 | cg01126010 | - | - |
| 27567156 | cg15843044 | cg15843044 | - | - |
| 27569287 | cg01861827 | cg01861827 | - | - |
| 27571486 | cg13958452 | cg13958452 | cg13958452 | - |
| 27571819 | cg13533303 | - | - | - |
| 27572615 | cg13533310 | - | - | - |
| 27572969 | cg13533317 | - | - | - |
| 27573249 | cg13533352 | - | - | - |
| 27573281 | cg13533354 | - | - | - |
| 27573369 | cg13533361 | - | - | - |
| 27573379 | cg13533362 | - | - | - |
| 27573534 | cg01589155 | cg01589155 | - | - |
| 27573550 | cg05151778 | cg05151778 | - | - |
| 27573652 | cg05990720 | cg05990720 | cg05990720 | - |
| 27573819 | cg23074747 | cg23074747 | cg23074747 | cg23074747 |
| 27573885 | cg13533397 | - | - | - |
| 27573891 | cg11613875 | cg11613875 | cg11613875 | cg11613875 |
| 27573968 | cg03854581 | cg03854581 | - | - |
| 27573982 | cg13533404 | - | - | - |
| 27573988 | cg14363787 | cg14363787 | cg14363787 | - |
| 27574186 | cg13533410 | - | - | - |
| 27574385 | cg01589701 | cg01589701 | - | - |

This table summarizes the CpG probes located within the *C9orf72* gene region, defined as the *C9orf72* gene and a 1 kb flanking region upstream and downstream, across various methylation platforms. The EPICv2 platform has 23 probes, the EPICv1 platform has 12 probes, the Methyl450K platform has 5 probes, and the Methyl27K platform has 2 probes.

### Replication in Independent Cohort

To assess whether these findings could be generalized to other cohorts, we validated our predictor in an independent DNA methylation cohort for the study of bipolar disorder, a sample that was readily available in our research group. This cohort includes 2,458 individuals (mean age of 51.1 [13.7], 57.3% female) without prior evidence of FTD or ALS. Briefly, this sample consists of n=1589 cases with bipolar disorder type I, n=289 independent controls, and n=580 first-degree relatives of the cases. The initial LASSO model from our prior analyses was applied to this independent cohort, and we identified four subjects with a predicted pathological *C9orf72* repeat expansion status. Two expansion carriers were diagnosed with bipolar disorder (one male and one female, both older than 50 years at the time of recruitment) and the other two were asymptomatic carriers (siblings of a bipolar disorder patient; both older than 50 years at the time of recruitment). We selected these four individuals and 17 randomly selected individuals of the cohort for independent experimental validation of their *C9orf72* repeat allele status by fluorescent repeat-primed PCR. The validation experiment was performed blindly to the predicted expansion status of these subjects. None of the 17 randomly selected individuals were identified by PCR as having pathological *C9orf72* repeat expansions, while all four predicted pathogenic repeat expansion carriers were indeed confirmed for having an expanded *C9orf72* repeat alleles in the pathogenic range (**Figure 3**).

**Figure 3.**
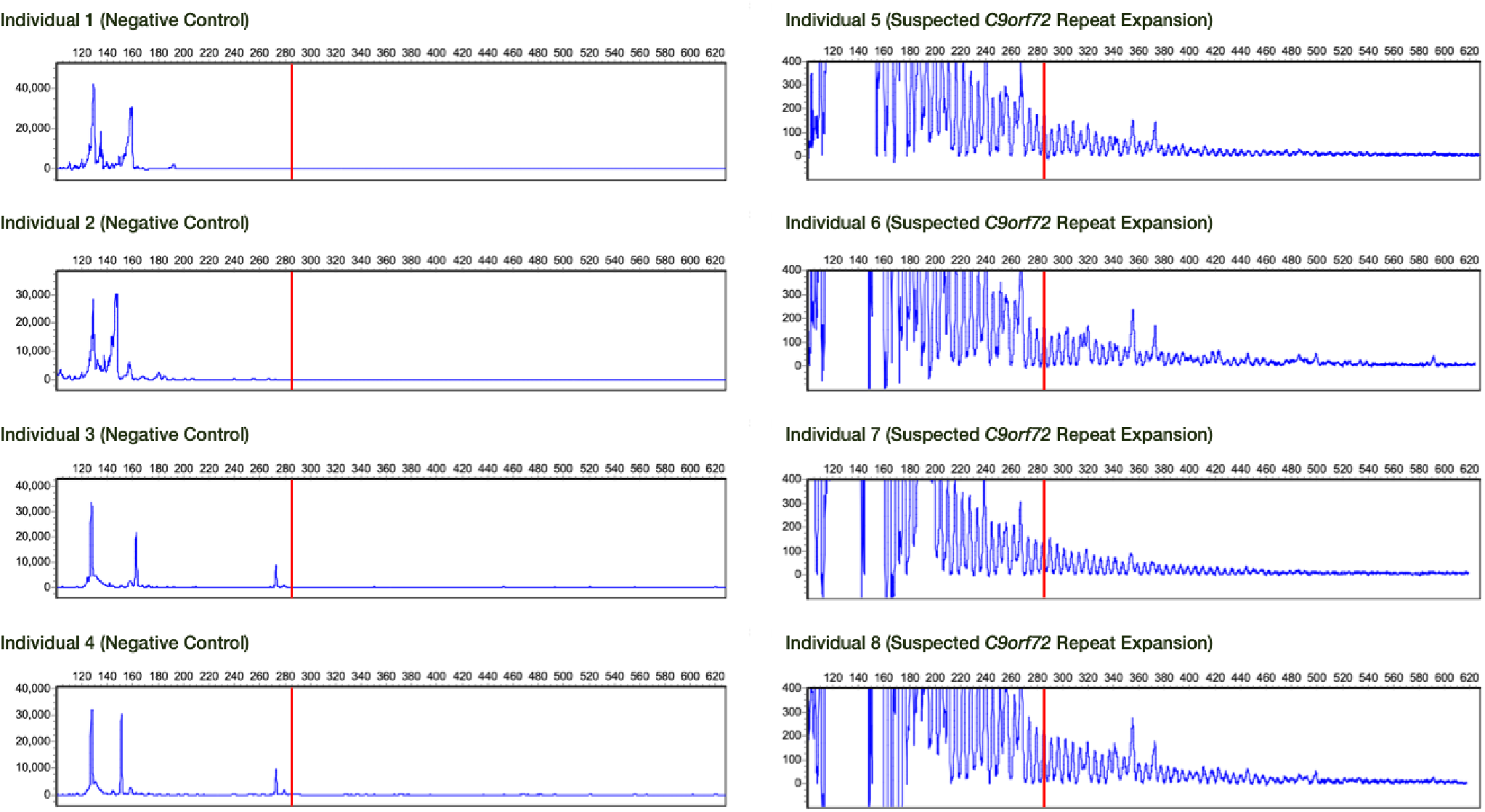
PCR Plots from Four Randomly Selected Negative Controls and Four Suspected Pathogenic *C9orf72* Repeat Expansion Carriers in the Replication Cohort. PCR Plots from four of the seventeen randomly selected negative controls from the validation cohort (Individuals 1-4) and four pathogenic *C9orf72* repeat expansion carriers identified by LASSO model (Individuals 5-8). The red line denotes the cut-off for what is considered to be a pathogenic repeat expansion length.

## Methods

### Cohort

Patients with frontotemporal dementia (FTD) (n = 318) and age- and sex-matched controls (n = 122) were included from the Amsterdam Dementia Cohort (ADC)^24^ for DNA methylation measurement from whole blood^24^. Six of the 440 samples were flagged as having a sample call rate error, and 157 individuals had no *C9orf72* repeat length data available, leaving 277 in the final test dataset (all FTD patients, mean age of 63.2 [8.3], 37.5% female). The ADC started collecting samples in the year 2000 and this is an ongoing, observational follow-up study of patients who visited the memory clinic of the Alzheimer Center at Amsterdam UMC, location VU University medical center (VUmc). All patients from the ADC underwent a standardized multidisciplinary assessment, consisting of medical history, informant-based history, neurological and medical examination, neuropsychological investigation, electroencephalography, brain magnetic resonance imaging (MRI), standard laboratory work-up and lumbar puncture. FTD was diagnosed according to diagnostic guidelines for FTD^25^. This study was performed in accordance with the ethical standards as laid down in the 1964 Declaration of Helsinki and its later amendments. Written informed consent was obtained from all participants.

### Repeat-primed PCR screening of the *C9orf72* repeat expansion

The measurement of the GGGGCC repeat of C9orf72 in the discovery cohort has been described in detail elsewhere^26^. PCR validation in the replication cohort was performed as described before^27^. In short, we used both fluorescent and repeat-primed PCR. Fragment length analysis was performed on an ABI 3730/ABI3730xI/3500 genetic analyzer (Applied Biosystems Inc., Foster City, CA, USA), and data were analyzed using the GeneScan software (version 4/5, ABI) and Peak Scanner Software, including a positive control sample for reference.

### DNA methylation measurement and preprocessing

DNA was extracted from whole blood using standardized protocols. 500 ng of each DNA sample was bisulfite converted with an EZ DNA methylation kit (Zymo Research, California, USA) following the manufacturer’s protocol specifically for a downstream analysis with an Infinium Methylation Microarray. Genome-wide methylation profiles were obtained using an Infinium MethylEPIC v2 BeadChip Kit (Illumina, California, USA) according to the manufacturer’s protocol.

A similar preprocessing pipeline was used for both the test dataset and the replication dataset. All data processing was performed using R 4.0.3. In the test dataset with FTD cases, six samples were removed for failing various technical criteria such as poor bisulfite conversions and poor hybridization or for having low fluorescence intensities. In the validation dataset, six samples were removed for the same criteria.

In both datasets, the following probes were removed: probes with low detection p-values, probes on sex chromosomes, probes with SNPs ^28^ at the CG or single base extension position, cross-reactive probes ^29^, and probes flagged by Illumina as problematic in the EPICv2 platform. In both datasets, probe removal was evenly distributed across chromosomes (**Figure S5**).

After a stringent preprocessing procedure, 434 individuals with 837,312 CpG probes remained in the test dataset and 2,458 individuals with 735,712 CpG probes remained in the replication dataset.

Since there is no established consensus on using M-values or beta values for statistical analysis, we opted to use M-values for our analyses.

### Differential Methylation Analysis

A differential methylation analysis was conducted using *limma* to identify differentially methylated probes between patients with *C9orf72* repeat expansion and without. DNA methylation values for each CpG probe were regressed against repeat expansion status. Age, sex, DNA methylation derived cellular composition ^23^, DNA methylation derived smoking score^22^, and experimental batch were included as covariates for the linear modeling. Genome-wide significance was initially set at *p* = 9e-08 in accordance with the literature ^30^. However, given the uneven distribution of *C9orf72* pathogenic repeat expansion carriers (n = 27) compared to non-carriers (n = 250), we conducted a permutation analysis by randomly shuffling carrier status across individuals 100 times to determine the amount of noise that is present in this type of analysis. The most significantly differentially methylated probe in these 100 permutations was subsequently used as the genome-wide significance threshold (*p* = 4.4e-10) for this methylation dataset.

### Prediction

To predict pathogenic repeat expansions in the *C9orf72* region, we restricted our analysis to the 23 CpG probes located within the *C9orf72* gene as well as 1kb upstream and downstream of the gene. The data was randomly split into a training set and a test set. The training set comprised 70% of the total set and the test set comprised 30% of the total set. A logistic regression model with L1 regularization was fitted using 10-fold cross-validation using the *glmnet* package on the training data. An initial single model was built to assess accuracy and identify which CpGs were identified as most important. Predictions were made on the test set using the fitted model with the best lambda value. To assess prediction accuracy across different array technologies, we conducted prediction analyses with the EPICv2 CpGs that were present in EPICv1, Methyl450K, and Methyl27K. For each of the four platforms, we randomly split the data 100 times. Each iteration involved training a LASSO model on 70% of the data and evaluating its accuracy on the remaining 30% to account for prediction variability due to data splitting.

### Independent Validation

The initial model trained with CpGs found in EPICv2 was applied to the independent validation dataset. These suspected repeat expansion carriers and 17 random selected subjects were identified as positive and negative controls respectively.

## Discussion

We performed a genome-wide differential DNA methylation analysis among carriers and non-carriers of pathogenic *C9orf72* repeat expansions in an FTD cohort. We identify eight CpG probes within the *C9orf72* gene region in pathogenic repeat expansion carriers that are uniquely and significantly hypermethylated after correction for multiple testing. Further analysis showed that DNA methylation data as obtained from the EPIC version 2 array (EPICv2) can be used to accurately predict individual carrier status of *C9orf72* repeat in the pathogenic range. When applied to an independent DNA methylation data set of roughly 2,500 subjects without diagnosis of ALS or FTD, the predictive model identified four carriers whose findings were validated by repeat-primed PCR. We observed an accurate prediction for *C9orf72* repeat expansion alleles in the pathogenic range with DNA methylation analysis, which may serve as a biomarker for epigenetic epidemiological studies with potential clinical relevance for ALS, FTD, and related phenotypes.

Identifying hypermethylation in eight CpG probes in the *C9orf72* gene for pathogenic repeat expansions is consistent with previous studies of ALS and FTD, identifying hypermethylation in the gene promoter region as well as the repeat itself ^20,21^. Hypermethylation has been noted in other GC rich repeat expansions. For example, the CGG repeat expansion in the X-linked *FMR1* gene associated with Fragile X Syndrome has been linked with hypermethylation at the promoter region and epigenetic silencing ^31,32^. Additionally, the GGC repeat expansion in the *XLT1* gene associated with Baratela-Scott Syndrome exhibited hypermethylation in the gene’s promoter region ^33^. Our genome-wide DNA methylation analysis results align with the current understanding of hypermethylation around the *C9orf72* repeat expansion being a neuroprotective mechanism to silence pathogenic gene expression^34,35^. It may be that the degree of hypermethylation of this locus in whole blood is clinically relevant for ALS/FTD disease status or progression, but further study is required to examine this in more detail. However, the primary effect of ALS/FTD pathology is neuronal which may not be detectable in whole blood.

Extending upon our differential methylation findings, the CpG probes at the *C9orf72* locus effectively predict the presence of repeat expansion alleles in the pathogenic range. However, these results were not generalizable across currently available DNA methylation array platforms. Restricting the model to include CpGs only present on the EPIC version 1 (EPICv1) array did not affect accuracy but limiting CpGs to those only present in the previous Infinium array types, i.e., Methyl450K and Methyl27K, resulted in significantly diminished accuracy and elevated Type I and Type II error rates. These results indicate that predicting *C9orf72* carrier status only appears to be effective using CpGs present on the higher resolution EPIC generation of DNA methylation arrays. Therefore, we recommend this prediction tool be used in DNA methylation cohorts assayed using EPICv1 and EPICv2.

Our DNAm predictor identified *C9orf72* repeat expansion carriers in a separate independent cohort, a finding that was thoroughly validated using repeat primed PCR, a gold-standard in *C9orf72* repeat expansion screening. The second cohort is part of an ongoing epigenetic study of severe mental illness with bipolar disorder cases, controls and first-degree relatives. Of the four individuals identified as having a *C9orf72* repeat expansion in the pathogenic range, two were diagnosed with bipolar disorder and the other two were unaffected siblings of a bipolar patient. There is no clinical evidence of ALS or FTD diagnosis or family history among these subjects and in-person clinical re-evaluation of these individuals is not possible. The link between *C9orf72* repeat expansions and psychiatric symptoms, however, remains a topic of interest^9,36^ as are the potential pleiotropic effects of repeat expansions in the pathogenic range in non FTD or ALS patients. The ability to identify carriers in epigenetic epidemiological study cohorts provides new opportunities to examine this further. We observe that the model’s complete accuracy in an entirely different cohort of individuals indicates that the model is not overfitted to the initial dataset, strongly validates the robustness and reliability of the model, and paves the way for its integration into other cohorts and broader research.

In the broad context of epigenetics, there is a growing body of research examining the relationship between methylation and its relationship to complex disorders. For instance, one recent study utilized a genome-wide DNA methylation analysis to identify DNA methylation variants (also known as epi-variants) at a single locus for neurodevelopmental and congenital anomalies that fail to be explained by conventional genetic testing. This analysis identified epi-variants that were associated with trinucleotide repeat expansion disorders^37^. In another study, researchers found upstream methylation that correlated with the length of the GAA repeat expansion in the first intron of *FXN*, a repeat expansion that results in Friedreich ataxia. These methylation patterns enabled prediction of expression of *FXN* and clinical outcomes in patients ^38^. Our research adds to this growing body of literature in providing a DNA methylation-derived tool for predicting repeat expansion status. To our knowledge, our work is one of the first studies to create a tool to predict individual repeat expansion status.

Epigenetic biomarkers have vastly broadened our understanding of aging. Machine-learning algorithms have been employed to build epigenetic measures of aging using tens to hundreds CpGs across the genome called epigenetic clocks ^39,40^. Clocks have also been trained to reflect age-related covariates like mortality and morbidity ^41,42^. For example, GrimAge is an epigenetic clock that combines DNAm-derived values like plasma protein estimates, smoking, age, and sex to determine mortality risk ^43^. While these clocks have been effective in predicting a variety of age-related variables, predicting cognitive impairment remains a challenge for many current epigenetic clocks. Some newer clocks like DUNEDinPACE^44^ have shown associations with certain dementia-related metrics like Alzheimer’s disease (AD) screening tests and some cognitive tests, but there remains a significant need for reliable measures of biological aging related to dementia and neurological diseases^45^. Integrating DNAm derived repeat expansions known to cause neurological conditions may offer additional granularity that improves the performance of epigenetic clocks in relation to dementia and other neurological disorders. This pathogenic *C9orf72* repeat expansion predictor may be the first of multiple DNAm-based repeat expansion predictors that aid in improving the performance of these dementia/cognition-based epigenetic clocks.

Predicting pathogenic repeat expansions in the *C9orf72* gene can also offer significant utility in epidemiological studies. Patients with *C9orf72*-associated FTD can often present neuropsychiatric symptoms like mania and psychosis without a family history, sometimes occurring years prior to more typical FTD symptoms^46^. Given the phenotypic heterogeneity of FTD, awareness of individuals with *C9orf72* pathological repeat expansion a priori in any large methylation dataset with patients with neurobehavioral conditions is extremely valuable^47^. Accurately identifying patients with bvFTD in the early stages is crucial for future FTD clinical trials^10^. The identification of four carrier subjects in the second cohort of about 2,500 individuals, suggests that these repeat expansions may be present at a higher prevalence than previously appreciated. A recent large-scale whole-genome sequence study observed an increased frequency of repeat expansion mutations across different populations^48^, which is in line with our finding of *C9orf72*.

There are a few limitations to this study. First, our dataset is small, and there are far more individuals without, than subjects with the pathogenic repeat expansion. However, the strength of the differential methylation analysis significant hits, given the size of this cohort, shows how strong and clear the association between the eight *C9orf72* CpGs and pathogenic repeat length is. Second, these findings are limited to whole blood DNA samples, which may not reflect the brain epigenetic patterns implicated in neurological conditions. Differing observations regarding somatic heterogeneity, referring to variation of the *C9orf72* repeat expansion size within an individual between tissue types, have been made in prior studies. One study observed somatic heterogeneity in individuals with large *C9orf72* repeat expansions, while another study found that *C9orf72* repeat expansion is consistent between brain and blood in asymptomatic or affected pathogenic *C9orf72* repeat expansion carriers^17,20^. Third, both this cohort and the independent cohort used for validation contain patients of European ancestry. The large *C9orf72* repeat expansion haplotype does seem to have a common European founder, but additional research needs to be done to see if these results hold in patients of varying ethnic and ancestral backgrounds ^8,49^.

In summary, the epigenetic *C9orf72* repeat expansion predictor offers significant utility at an individual clinical level, at a population level for large DNAm cohorts, and at a research level, for study of neurodegenerative disorders, of pleiotropic effects of *C9orf72* repeat expansions, epigenetic aging, and neurological and cognitive outcomes. We expect that other repeat expansions and types of genomic variation are also detectable via epigenetic profiling and that this DNAm predictor will be a first among others to be used clinically and in research.

## Supplementary Materials

**Figure S1:**
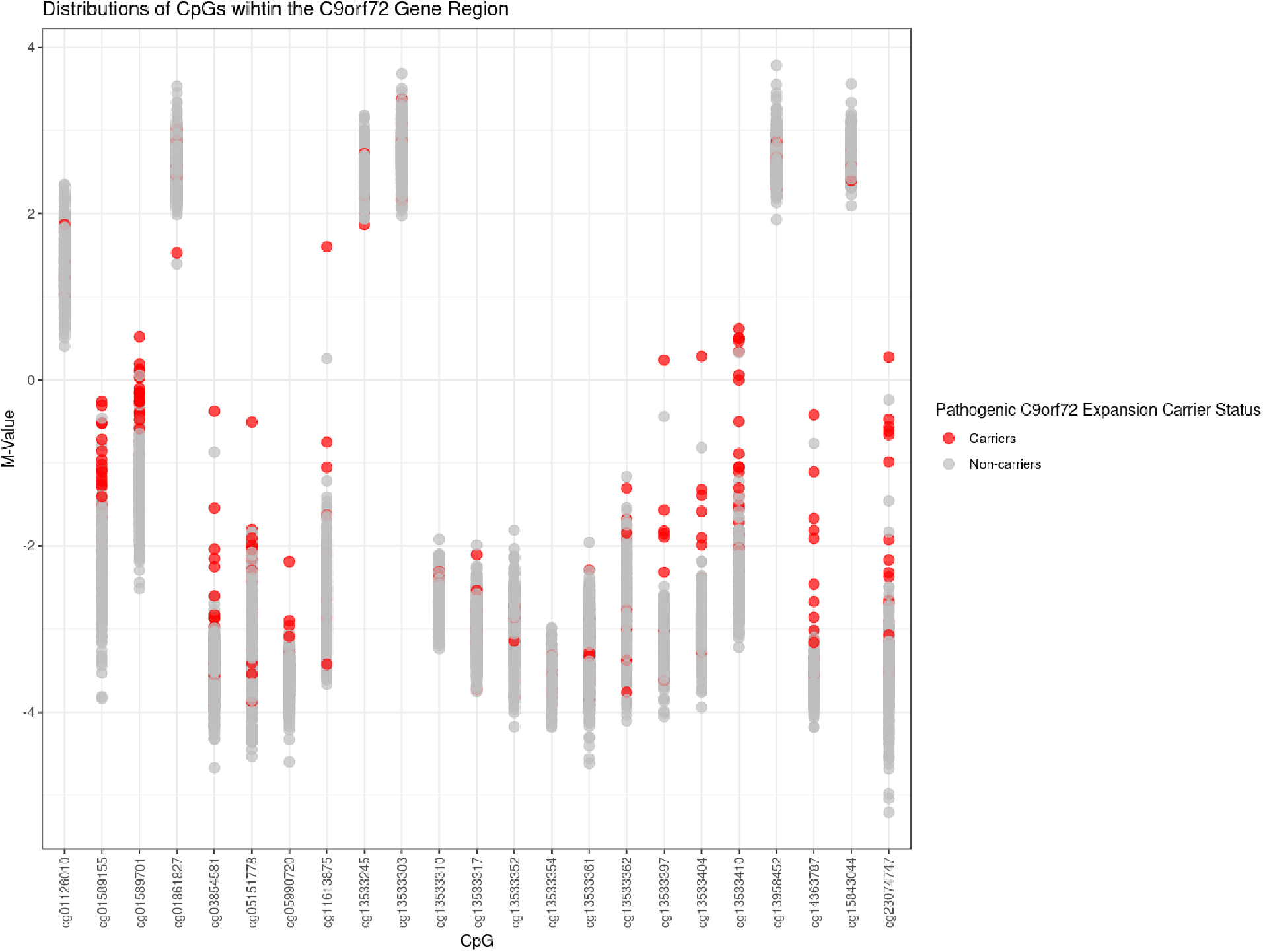
Distributions of CpGs within the C9orf72 Gene Region. This plot shows the distribution of M-Values of the CpGs in the *C9orf72* gene region (within the gene and 1 kb upstream/downstream). Individuals are colored red if they are carriers of a pathogenic *C9orf72* repeat expansion and colored grey if they are non-carriers.

**Table S1:**
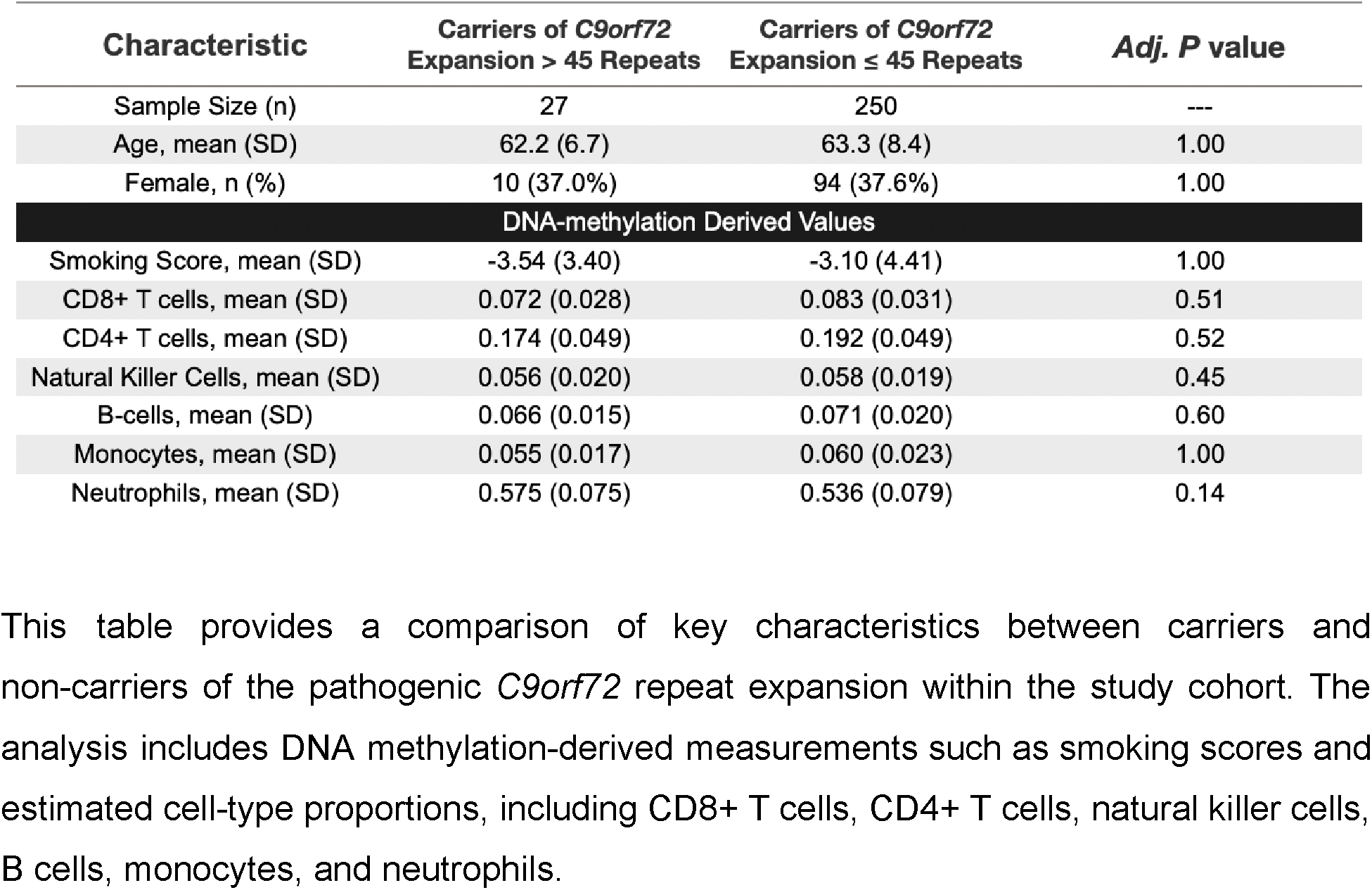
Sample Characteristics From Amsterdam Dementia Cohort (ADC)

| Characteristic | Carriers of *C9orf72*<br>Expansion > 45 Repeats | Carriers of *C9orf72*<br>Expansion ≤ 45 Repeats | Adj. *P* value |
| --- | --- | --- | --- |
| Sample Size (n) | 27 | 250 | --- |
| Age, mean (SD) | 62.2 (6.7) | 63.3 (8.4) | 1.00 |
| Female, n (%) | 10 (37.0%) | 94 (37.6%) | 1.00 |
| DNA-methylation Derived Values |  |  |  |
| Smoking Score, mean (SD) | -3.54 (3.40) | -3.10 (4.41) | 1.00 |
| CD8+ T cells, mean (SD) | 0.072 (0.028) | 0.083 (0.031) | 0.51 |
| CD4+ T cells, mean (SD) | 0.174 (0.049) | 0.192 (0.049) | 0.52 |
| Natural Killer Cells, mean (SD) | 0.056 (0.020) | 0.058 (0.019) | 0.45 |
| B-cells, mean (SD) | 0.066 (0.015) | 0.071 (0.020) | 0.60 |
| Monocytes, mean (SD) | 0.055 (0.017) | 0.060 (0.023) | 1.00 |
| Neutrophils, mean (SD) | 0.575 (0.075) | 0.536 (0.079) | 0.14 |

This table provides a comparison of key characteristics between carriers and non-carriers of the pathogenic *C9orf72* repeat expansion within the study cohort. The analysis includes DNA methylation-derived measurements such as smoking scores and estimated cell-type proportions, including CD8+ T cells, CD4+ T cells, natural killer cells, B cells, monocytes, and neutrophils.

**Figure S2:**
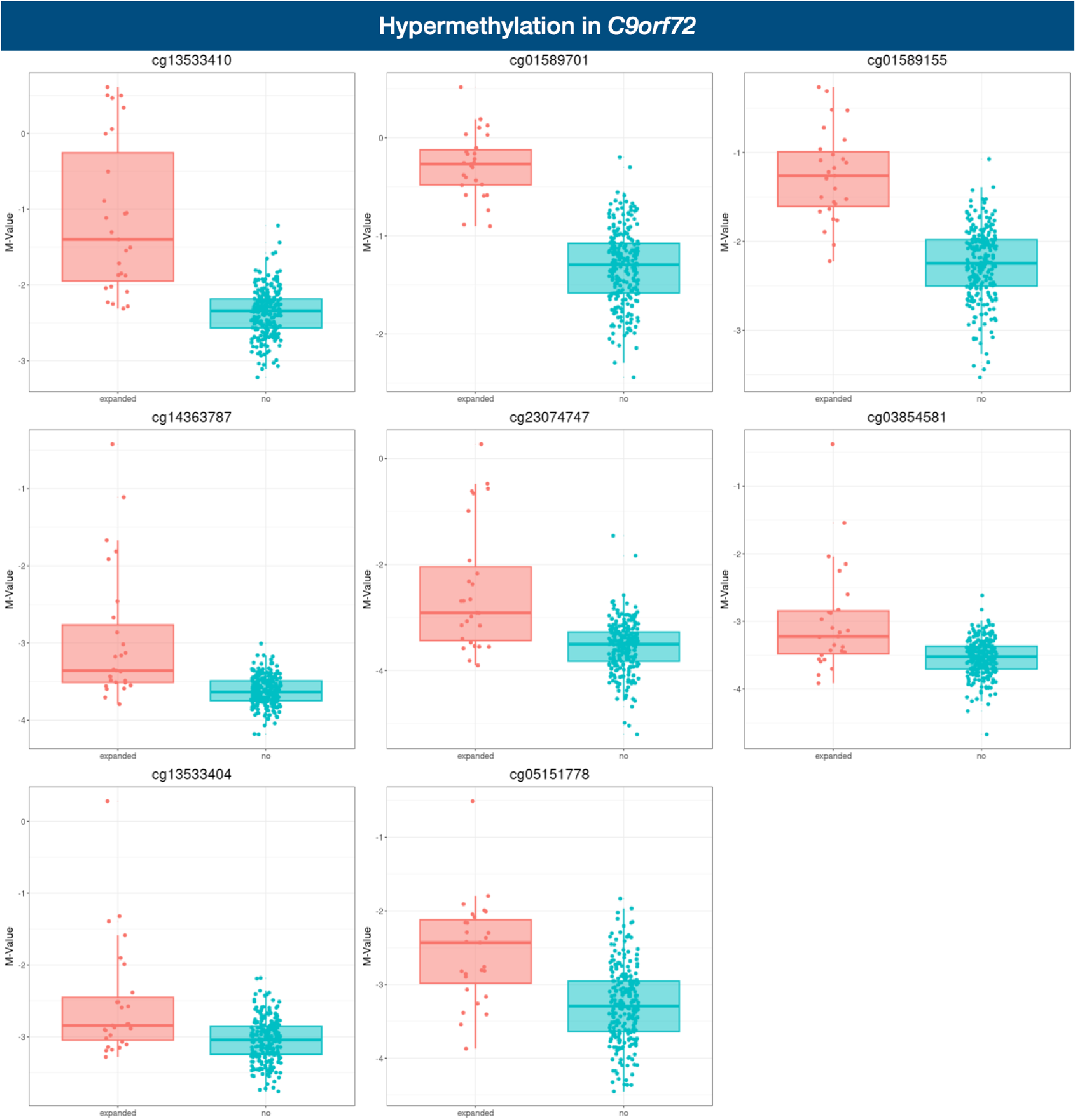
Genome-Wide Significant Differential Methylation Between *C9orf72* Pathogenic Repeat Expansion Carriers and Noncarriers. This figure shows the distribution of M values for the eight CpG sites identified as significantly differentially methylated between carriers and non-carriers of the pathogenic *C9orf72* repeat expansion.

**Figure S3:**
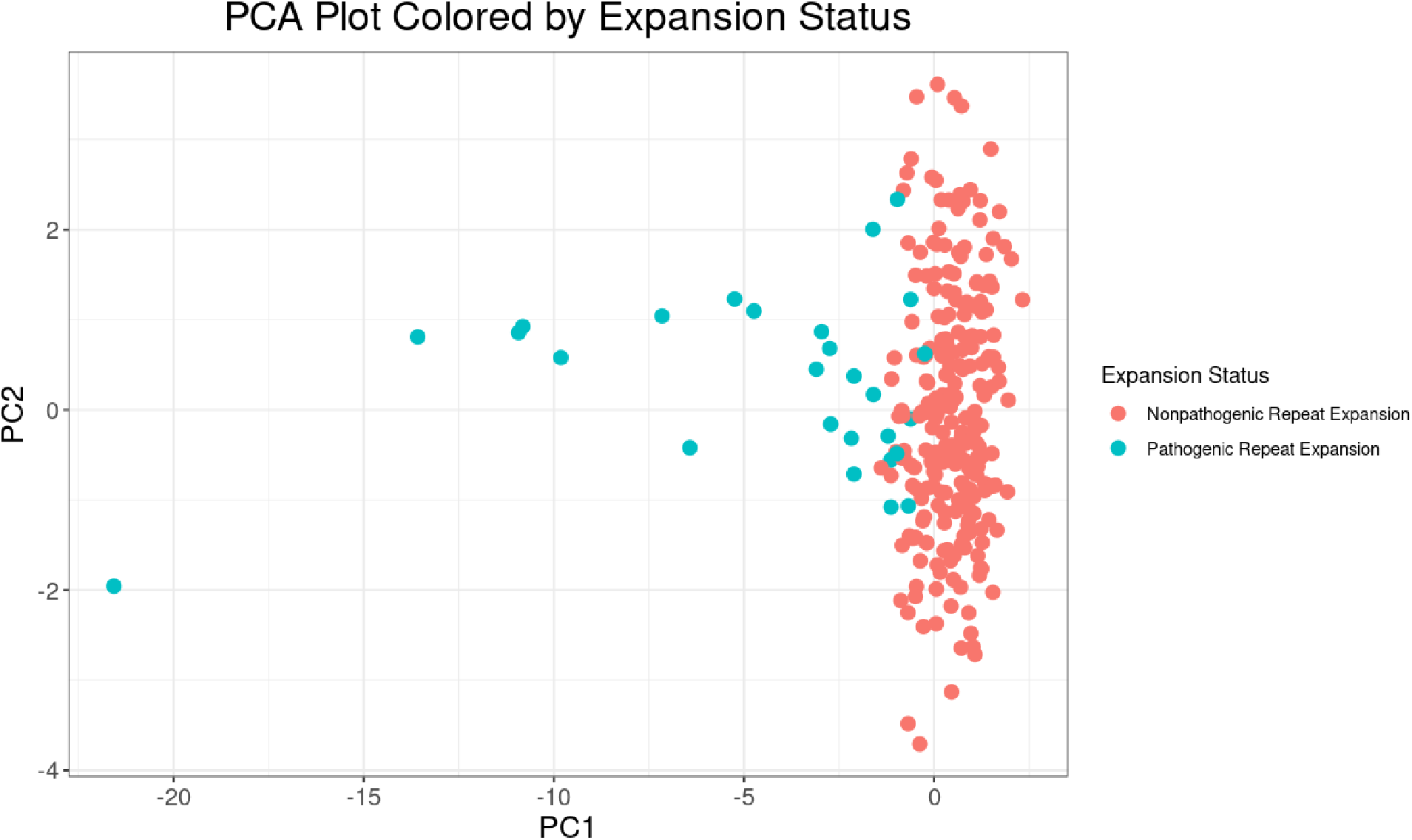
Principal Component 1 vs. 2 of 23 CpG Probes in *C9orf72* Gene Region. This PCA plot compares principal components (PC) 1 and 2 of the CpG probes in the *C9orf72* gene region, colored by whether an individual was a carrier or non-carrier of a pathological *C9orf72* repeat expansion. There is evident separation between individuals with and without pathological *C9orf72* repeat lengths.

**Figure S4:**
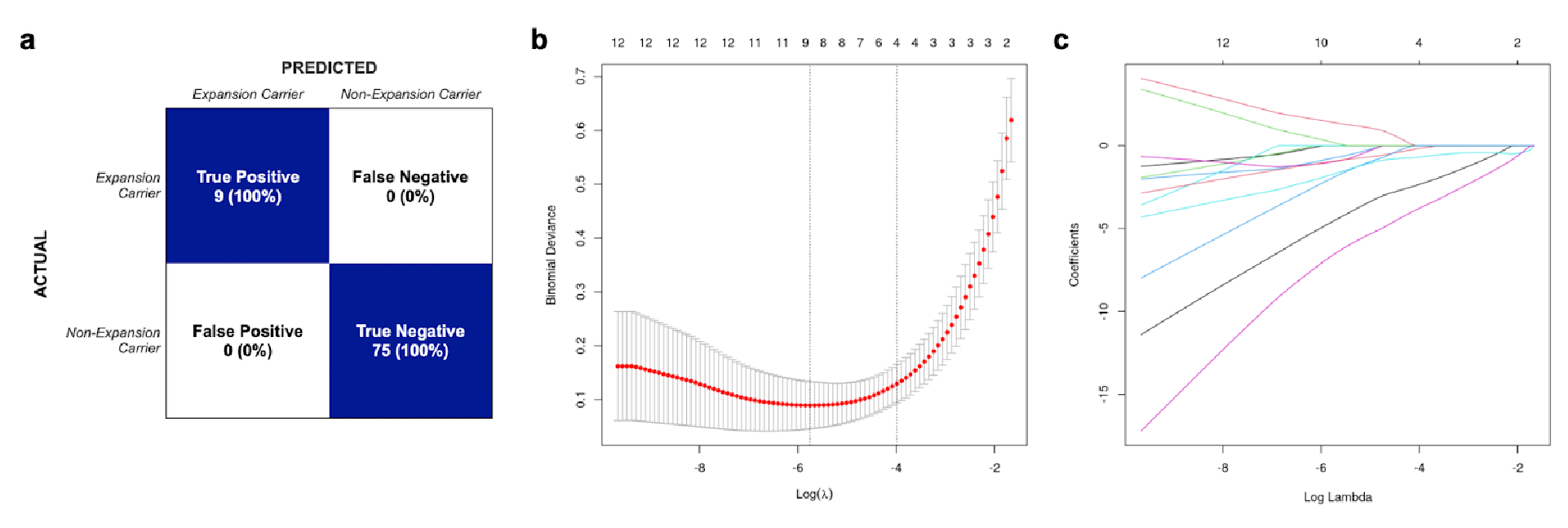
Prediction Confidence Matrix and LASSO Regression Variable Selection. This figure shows the application of Least Absolute Shrinkage and Selection Operator (LASSO) regression to predict pathological *C9orf72* repeat expansion status based on methylation profiles of the 23 CpG probes in the *C9orf72* gene region. (a) Confusion matrix showing 100% accuracy in predicting carrier status in the test set. (b) Cross-validation plot optimizing the lambda parameter to minimize error. (c) Coefficient profiles of the 23 CpG probes as a function of lambda.

**Table S2:**
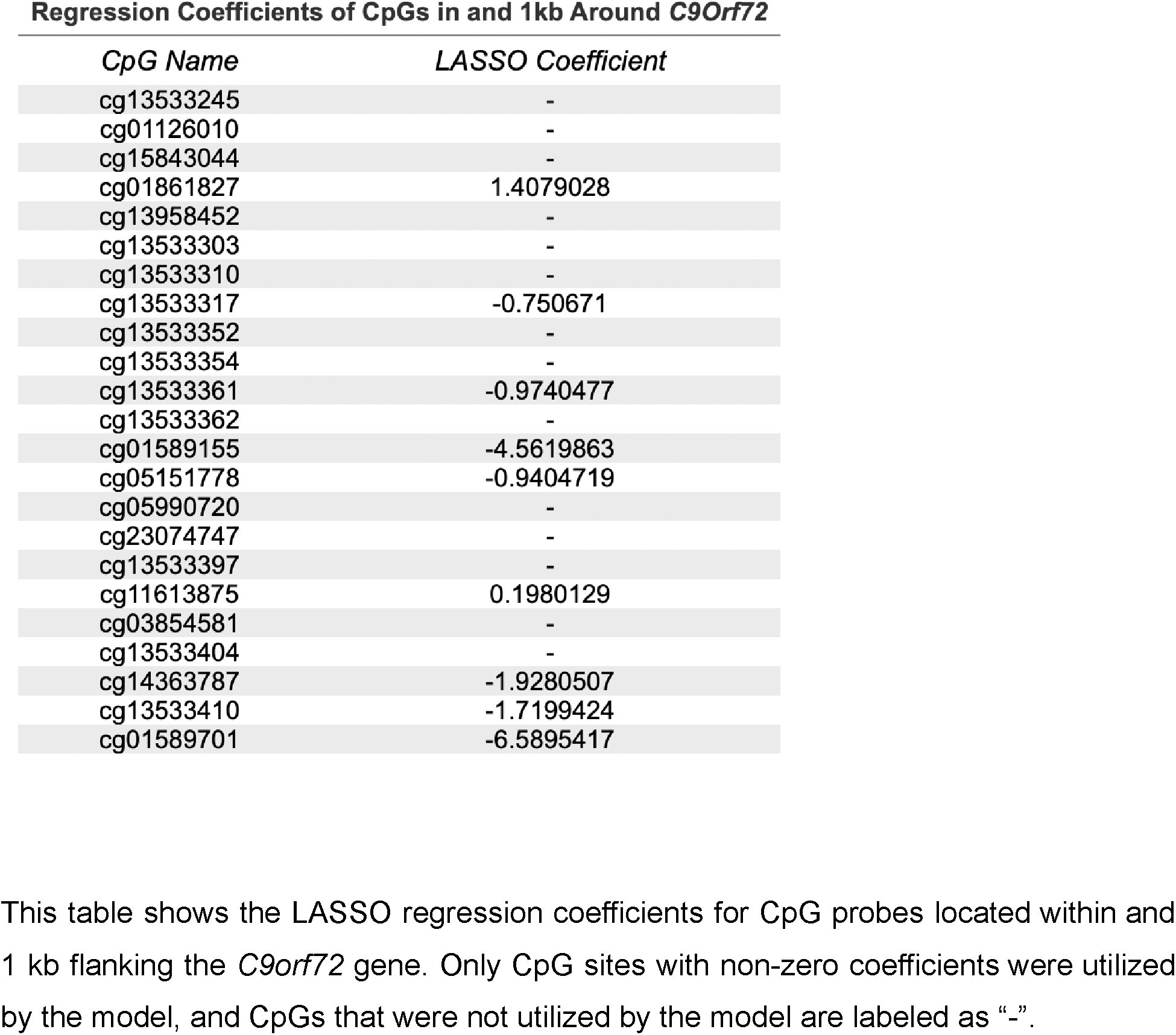
LASSO Regression Coefficients of CpGs in *C9orf72* Gene Region in Initial Model.

**Figure S5:**
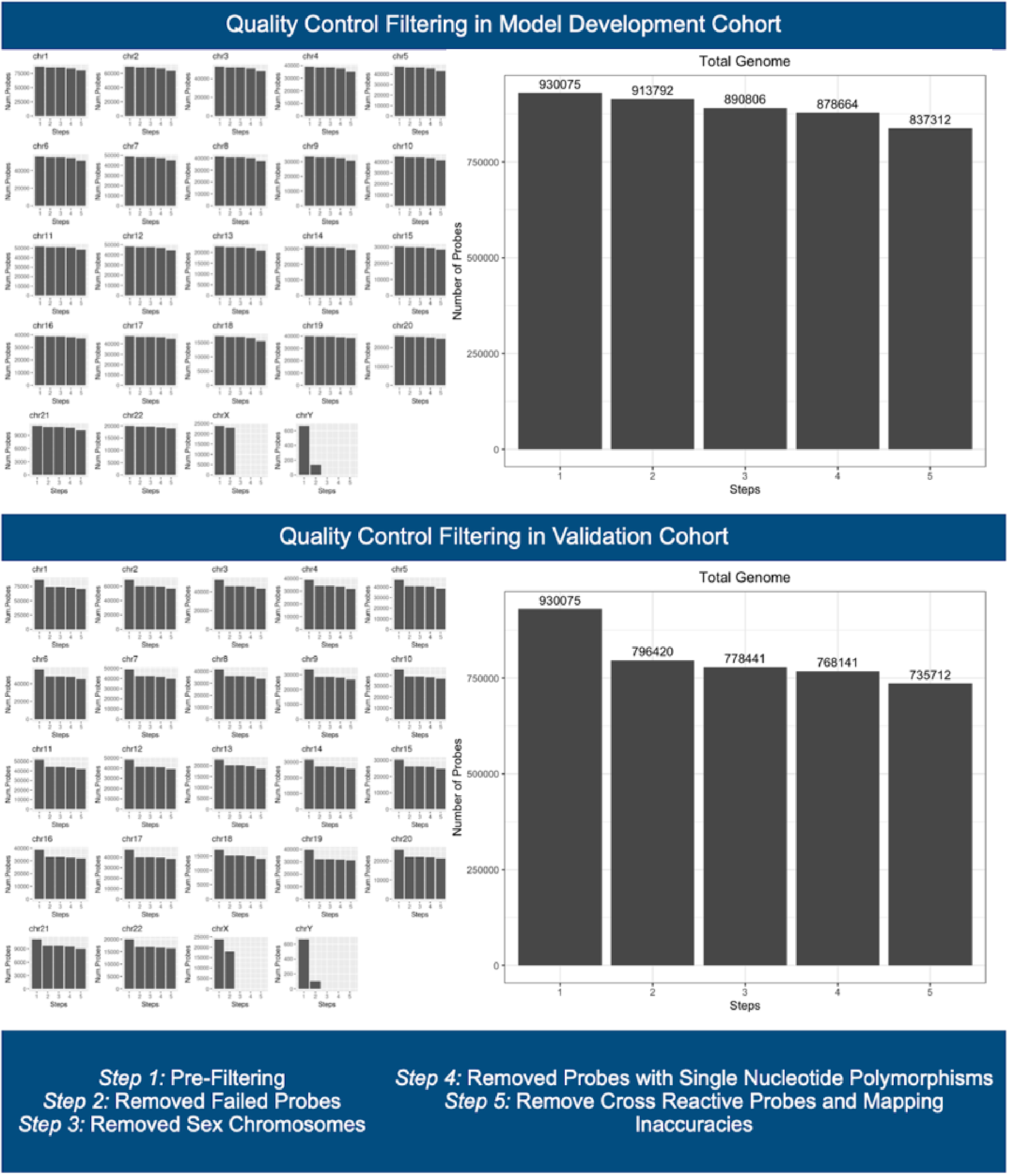
Quality Control Filtering in Both Cohorts. This figure displays the quality control filtering applied to the methylation data in both the model development and validation cohorts. In the model development cohort, probes (930,075) were filtered through five steps: (1) pre-filtering, (2) removal of failed probes, (3) exclusion of sex chromosome probes, (4) exclusion of probes with single nucleotide polymorphisms (SNPs), and (5) removal of cross-reactive probes and mapping inaccuracies, leaving 837,312 probes. In the validation cohort, similar filtering steps were applied, resulting in 735,712 retained probes after quality control. In both cohorts, each quality control step led to a fairly even distribution of filtering across chromosomes.

## Acknowledgments

Research of Alzheimer center Amsterdam is part of the neurodegeneration research program of Amsterdam Neuroscience. Alzheimer Center Amsterdam is supported by Stichting Alzheimer Nederland and Stichting Steun Alzheimercentrum Amsterdam. The clinical database structure was developed with funding from Stichting Dioraphte. We are greatly appreciative to those individuals who donated blood samples on which this study was based. LMR was funded by the Memorabel fellowship (ZonMW projectnumber: 10510022110012). The FTD DNA methylation study was supported by the National Institute on Aging (NIH/NIA) Grant No. R21 AG072390.

